# Not so selfish: transposable elements fuel adaptation to climate change

**DOI:** 10.1101/2024.09.02.610836

**Authors:** Evgenii V. Potapenko, David Schwarts, Tali Mandel, Nimrod Ashkenazy, Dana Fuerst, Guy Atsmon, Abraham B. Korol, Michael B. Kantar, Avi Bar-Massada, Sariel Hübner

**Affiliations:** Department of Bioinformatics and Galilee Research Institute (MIGAL), Tel Hai College, Upper Galilee, 12210 Israel; Department of Evolutionary and Environmental Biology, University of Haifa; Haifa, Israel; Department of Tropical Plant and Soil Sciences, University of Hawai’i at Mânoa, Honolulu, HI, USA; Faculty of Natural Sciences, University of Haifa, Kiryat Tivon 36006, Israel

## Abstract

Biodiversity conservation is urged at biodiversity hotspots that are under constant threat from anthropogenic development, yet a careful examination of the adaptive potential is a prerequisite for action. The Levant is considered a biodiversity hotspot and the distribution edge for many species, including important crop wild relatives. This region is under accelerated desertification and constantly disturbed by human activities, thus urging intervenient action. We collected and sequenced 300 wild barley plants along an eco-geographic gradient following a unique ecological-genetic sampling design. This scheme enabled to overcome the tight correlation between environmental and geographic distances. Phenotypic data was collected from 3600 progeny plants over three years and enabled to identify adaptive haplotype blocks comprised of phenological regulating genes tightly linked to drought and heat responsive genes. These haplotype blocks were highly enriched for transposable elements insertions, likely regulating genetic variation around adaptive genes, especially in stressed populations. Ecological and evolutionary models using over 2600 observations were combined to predict maladaptive risk, indicating that populations will be funneled into higher water availability refugia habitats while increasing isolation. Our findings highlight the main factors affecting rapid local adaptation and provide important recommendations for biodiversity management and conservation.

## Introduction

Accelerated climate change and destruction of natural habitats, specifically in the past century, are threatening biodiversity, agriculture, and human health worldwide^1,2^. To pace with the projected environmental change, species must rapidly evolve through genetic adaptation. Adaptation to a changing environment is derived through a dynamic venture of interacting evolutionary forces, which can act gradually on a large panel of small contributing alleles^3^ or a few large-effect loci^4^. Apart of the strength and type of selection, the pace and direction of the subsequent evolutionary change is determined by other genetic and demographic factors including linkage disequilibrium, migration, and drift^5^. Intuitively, local adaptation at heterogenous environment evolves through divergent selection favoring altered beneficial genetic variation at different habitats while overcoming contrasting forces of gene flow and drift^6,7^. Theory predicts that local adaptation can be accelerated by favoring the evolution of linkage between beneficial loci^8–10^. Thus, adaptive response to environmental change can be accelerated when interactions evolve into a concentrated heritable genomic architecture which harbors several beneficial alleles^11,12^. Other modes of accelerated local adaptation are attributed to specific events of chromosomal modifications like inversions and deletions^13,14^, yet the association with gradual or fluctuating environmental change remains elusive. A key dynamic portion of the eukaryotic genome is comprised of transposable elements which are highly abundant in plants^15^, and are activated in response to environmental stress. These elements can regulate the expression of adaptive genes and lead to altered chromosomal organization, thus providing a mechanism of rapid response to environmental change^16,17^.

The accessibility of genomic data enables the development of predictive models for populations dynamics from an evolutionary perspective by estimating the adaptive potential to future climate^18–20^. This intriguing approach is attractive, yet the high rate of false positive signals impedes its predictive power^21^. To gain a more nuanced understanding of risks induced by climate change, it is imperative to integrate ecological and genomic models to assess future habitat suitability in the context of evolving populations^22–24^. However, the complex interplay between natural selection, demographic factors, and the environment frequently leads to the identification of spurious genomic signals of adaptation with vague biological interpretation. To overcome these caveats, a careful and targeted sampling design must be applied to avoid some of the inherent correlations between genetic and geographic distances^25^. Moreover, a phenotypic evaluation of the studied populations can potentially improve the interpretation and reliability of the identified adaptive signals and consequently the power of predictive models^17,26^. The impact of climate change is specifically alarming in transitional regions that are rich in biodiversity, including wild relatives of many crops which are an important source of beneficial alleles for breeding^27^. The Eastern Mediterranean basin (Levant) bridges between Africa and Eurasia, and is considered as a global biodiversity hotspot and home to many crop wild relatives^28,29^. The dense human population and intensive infrastructure development in this region call for the adoption of knowledge-based conservation strategies along with cautious interference and management practices. Wild barley (*Hordeum spontaneum*) populations occur in large stands along a wide range of habitats in the southern Levant, from the Mediterranean timberline zone at high elevation (1600m above sea level) to desert climate near the Dead Sea (−400m), thus providing an excellent system to study local adaptation^30^. However, the geographical setting induces a tight link between latitude and environmental clines (north-Mediterranean, south-desert). This correlation constrains the identification of genetic factors that underlie local adaptation. Moreover, the high rate of self-fertilization in wild barley^31^ further increases the impact of isolation by distance on the landscape of genetic diversity. Previous collections of wild barley in this region^30,32^ were not designed to tackle these issues and thus are less suitable for identification of genome-environment associations and development of predictive evolutionary models.

Here, we describe the establishment of a wild barley collection designed to study local adaptation. Whole genome sequence data was generated for all accessions and used to identify the genetic mechanisms involved in rapid local adaptation. Understanding these mechanisms and identifying the underlying genes enabled the development of conclusive predictive models for risk assessment which can inform conservation and management strategies to mitigate the adverse effects of climate change.

## Results

### Local adaptation is derived by adaptive genes found in TE-rich regions

We sampled 300 individual plants from 30 different sites along the southern Levant following an ecological-genetic sampling design to overcome the tight correlation between adaptive and demographic factors (Fig. S1). Thus, we targeted for sampling populations that occur at similar environmental conditions (e.g., precipitation, temperature, soil type) but are genetically assigned to distinct genetic clusters^30,33^ at different geographical regions (Fig. 1a). To evaluate population structure and potential admixture, data was pruned to reduce bias imposed by tight linkage disequilibrium (LD), leaving a total of 879,897 SNPs for PCA and structure analyses (Fig. 1b). Wild barley populations were largely assigned to clusters following the expected split to ecotypes with a gradual transition between ecotypes (Fig. 1b). In accordance with our hypothesis, the sampling design indeed reduced the correlation (*r_Mantel_* = −0.33, *p* = 0.98) between geographic and genetic distances (Fig. S2) compared to previous collections in this region^30^.

**Figure 1.**
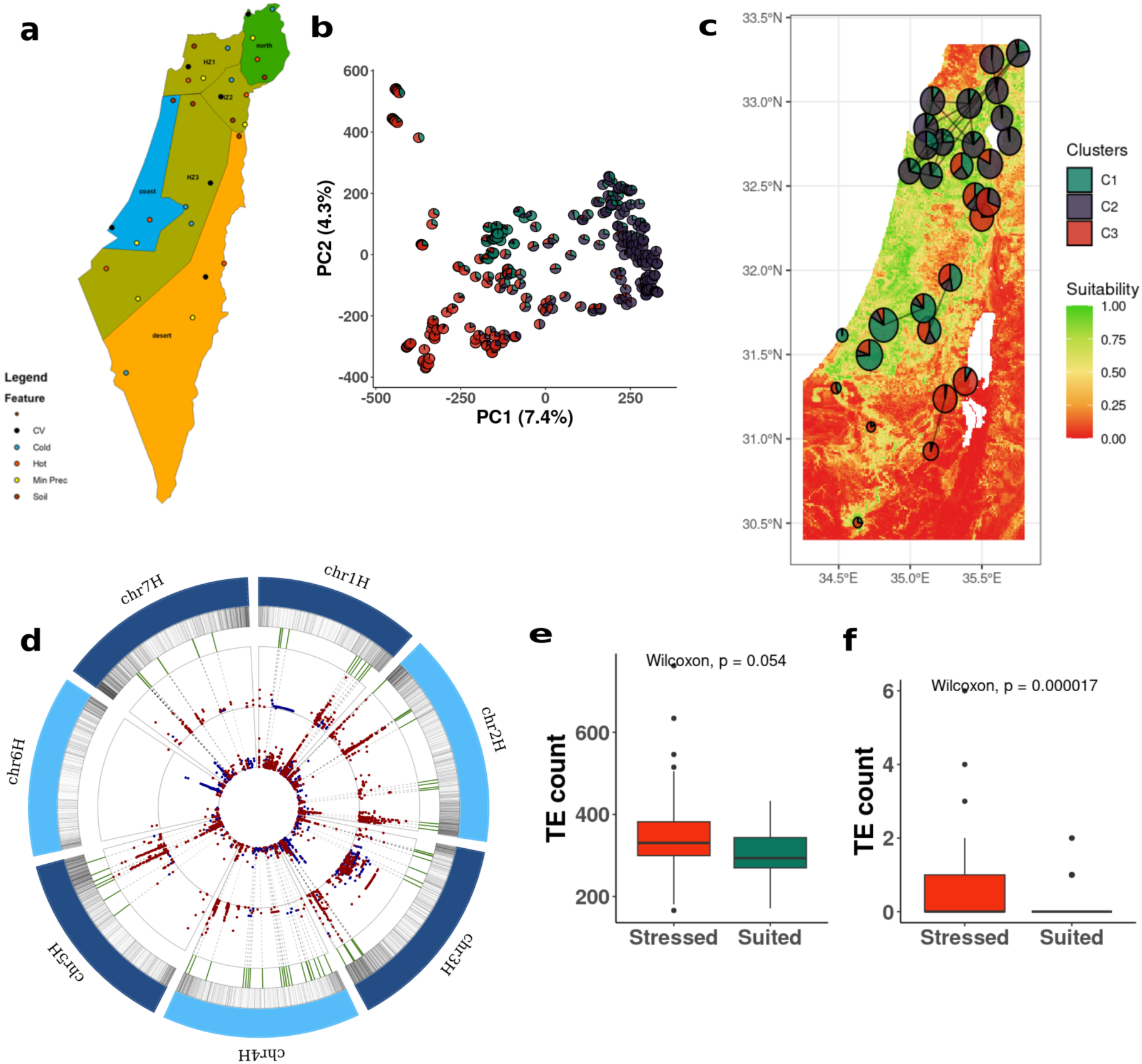
Exploring genomic regions of barley associated with local adaptation. **a)** A map depicting the sampling design. The geographic area was split into six polygons in accordance with ecotypes boundaries, genetic groups, and ecotones. Within each polygon, 5 sampling locations were targeted based on ecological conditions. **b)** PCA of all 300 accessions with pie charts representing assignment to the three genetic clusters identified by structure analysis. **c)** Species distribution model for current environmental conditions where the color gradient on the map depicts the suitability score, the pie charts colors illustrate assignment to genetic clusters, and their size indicate Tajima’s D statistics (smaller pie correspond to lower score). Black lines between populations represent gene flow. **d)** Circular plot of adaptive HBs. From outer circle inwards: transposable elements insertions density, adaptive HBs indicated by green vertical lines significant associations with climatic factors based on EMMAX by LFMM. SNPs associated with temperature factors are marked in red, and precipitation in blue. Dashed lines highlight SNPs within overlapping HBs. **e)** Comparison of TEs insertions between stressed and non-stressed populations. **f)** Comparison of TEs insertions around adaptive regions.

To identify genomic regions that contribute to local adaptation along environmental clines genome-environment association (GEA) analyses were performed using the seven most relevant bioclimatic variables, namely annual mean temperature (BIO-1), mean diurnal range (BIO-2), isothermality (BIO-3), temperature seasonality (BIO-4), temperature annual range (BIO-7), annual precipitation (BIO-12), and precipitation seasonality (BIO-15) from WorldClim2 database^33^. To improve the association signal, we calculated the identity-by-state (IBS) across all accessions and filtered-out samples with tight genetic similarity leaving 218 accessions for the GEA analyses. The SNP dataset was filtered accordingly to capture the changes in minor allele frequency and missing data, leaving a total of 27,239,486 SNPs for the GEA analyses. We used three different GEA approaches (EMMAX^35^, LFMM^36^ and RDA^37^) and overlapped the significant associations obtained in each analysis after Bonferroni correction to robustly identify adaptive loci. The RDA analyses yielded high proportion of significant signals compared to other analyses, likely due to higher false positive rate (Fig. S3). Thus, we considered a candidate adaptive region only when supported by both LFMM and EMMAX (Fig. 1d; Fig. S4). Altogether, 894 significant SNPs were identified within 113 independent haplotype blocks (HBs) that were evenly spread along chromosomes. Among HBs, 95 blocks were longer than 10Kbp of which 74 HBs were associated with temperature related bioclimatic factors, 16 HBs with precipitation factors and 5 HBs were simultaneously associated with both precipitation and temperature (Table S1).

The concentrated genetic architecture within HBs can potentially facilitate a synchronized adaptive response to environmental change, yet a rapid mechanism for inducing genetic variation at those regions is necessary in order to maintain high adaptability over time. We hypothesized that the activation of transposable elements (TEs) under environmental stress can potentially serve as an inducing mechanism for genetic variation around adaptive HBs. To test this, we compared the accumulation of TEs insertions in stressed versus non-stressed populations. To identify populations that are currently under environmental stress, we calculated the habitat suitability for wild barley using a random forest species distribution model (SDM) (Fig. S5). To improve the model performance, the number of observations was expanded to include 2641 locations where wild barley populations were recorded in the last decade. High prediction accuracy (AUC > 0.92) was obtained for the models under different scenarios, and the distribution of wild barley and suitability of habitats well supported *in-situ* observations of population densities, thus higher suitability under Mediterranean conditions than in the desert (Fig. 1c). Based on these models, we inspected the profile of TEs insertions along the genomes in accessions that were sampled from habitats of highest and lowest suitability (Fig. 1d). Accessions from less suitable habitats accumulated across the entire genome mildly more TEs than those from highly suitable habitats (*W =* 774*, p =* 0.053, Fig. 1e). However, focusing on the adaptive genomic regions identified by the GEA analyses, a significantly higher rate of TEs insertions were observed in accessions from low suitability habitats (*W* = 9134, *p* = 0.000017, Fig. 1f). To confirm this trend, the extent of insertions surrounding adaptive regions was compared to a set of regions sampled randomly along the genome (excluding pericentromeric regions to avoid bias). Again, a significantly higher rate of insertions was observed around adaptive regions than expected by chance (*n* = 1000, *p* = 0.007). These results indicate the involvement of TEs accumulation around adaptive haplotype blocks, presumably contributing to generation and maintenance of genetic variation.

### Climate change increases geographic and phenological isolation between populations

To predict current and future (2070) habitat suitability for wild barley, SDM were ran for different scenarios. In the models, four static soil and geographic parameters (pH, cation exchange capacity, distance to river, elevation) were included together with three predicted bioclimatic factors (BIO-2, BIO-15, BIO-16). Collinearity between variables was controlled and erratic parameters (e.g. NVDI, vegetation) which cannot be robustly predicted in the future were excluded. Highest habitat suitability under current conditions was obtained at lower elevations along the Mediterranean region and at high elevations in the desert, where precipitation is the main limiting factor (Fig. 1e, 2a,b; Fig. S5). Under the considered scenarios for future climate, a maximum reduction of 7% in suitability was predicted across habitats. Reduction in suitability was pronounced in the drier regions (annual precipitation < 300mm) along the Jordan valley in the east, and the Negev desert in the south (Fig. 2a). Moreover, the expected shift in precipitation was also reflected with a reduction in suitability along the southern coastal region (Fig. 2a). The pattern of declining suitability manifested under mild and severe scenarios counterparts the ongoing process of desertification across the Levant. Despite the predicted loss of habitats along this region, high suitability will be maintained along creeks of large drainage basins, and along mountain ridges where temperatures are expected to remain mild also under future conditions (Fig. 2a). The improvement in suitability along creeks and higher elevations is already noticed today in the desert and is expected to expand in the future to other regions. Desert populations are more isolated (Average IBD = 0.7) compared to Mediterranean populations (Average IBD = 0.47; *t* = 3.54, *p* = 9.3×10^-4^) which is also reflected with a 35% reduction in nucleotide diversity in the desert (Fig. S6). The lower diversity and reduced gene-flow among desert populations are likely to further constrain their ability to maintain the necessary genetic variation to adapt and sustain future climate. This trend is likely to increase isolation among populations, generating refugia habitats for wild barley and likely also for other species.

**Figure 2:**
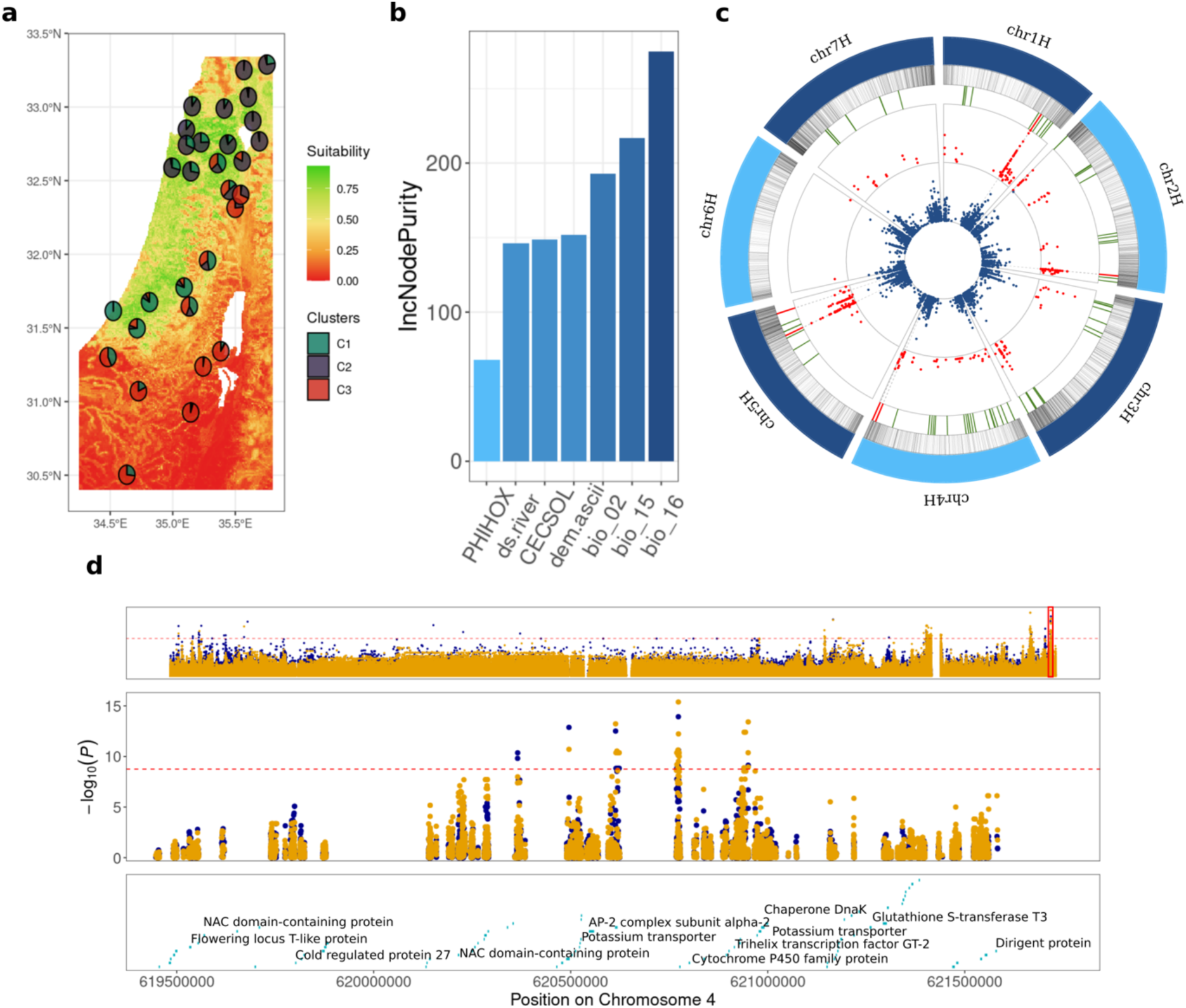
Genomic footprints of local adaptation under climate change. **a)** Species distribution model predicted for future (2070) climate, where the color gradient on the map depicts the suitability score. **b)** Relative contribution of the environmental factors used in the predicted species distribution including pH (PHIHOX), distance from river (ds.river), cation exchange capacity of the soil (CECSOL), elevation (dem.ascii), mean temperature range (BIO-2), precipitation seasonality (BIO-15), mean precipitation in winter (BIO-16). **c)** Circular plot of adaptive genomic regions. From outer circle inwards: transposable element insertions density, locations of adaptive HBs (green) and the focal six regions (red), overlapping significant associations in the GEA analyses (red dots), associations with the flowering time (blue dots); dashed lines indicate overlaps between GEA and GWAS for flowering time. **d)** Detailed view of an adaptive haplotype block on chromosome 4 showing significant associations with BIO-3 as detected by EMMAX (**blue**) and LFMM (**orange**). At the top panel the entire chromosome is plotted with the marked region in red rectangle. The middle panel is a zoom-in to the highly significant region where horizontal red dashed line corresponds to Bonferroni correction and the candidate adaptive genes are indicated at the bottom panel.

To evaluate the adaptive phenotypic response to climate change, a common garden experiment was performed. A total of 3600 plants were grown and screened for key phenological and morphological traits over three years. A wide range of phenotypic features was recorded in accordance with the environmental variation along habitats (Table S2). For example, flowering time extended over 74 days between the first flowering desert accessions (73 days from sowing) to the late flowering accessions from Mt. Hermon (147 days from sowing). A similar trend was also observed in morphological traits such as spike length, where spikes were four times longer in northern accessions (max = 22cm) compared with desert ecotype (min = 5cm). Among the measured traits, heritability was highest for flowering time (*H^2^* = 0.57) and lowest for number or tillers (*H^2^* = 0.12), implying that the adaptive response is attributed to synchronized flowering (escape strategy) while reduction in biomass is a plastic feature which is more responsive to changes in environmental conditions between years. To identify genomic regions and candidate genes that underlie the measured traits, we performed GWAS corrected for population structure and relatedness. To link the results with adaptation along environmental gradient, the identified top signals in the GWAS were overlapped with the HBs identified in the GEA analyses (Fig. 2c, Fig. S7, Table S1). Interestingly, all overlapping signals were supported by two or more phenotypic traits suggesting a pleiotropic regulation of adaptation and enhanced adaptive response projected by a clustered genetic architecture. These regions were further explored for potential candidate adaptive genes that are affected by environmental signal and contribute to functional phenotypic response (Fig. 2d, Table S3). All inspected genomic regions of adaptive response harbored clusters of tightly linked genes that are associated with response to abiotic stress. For example, a strong overlapping signal associated with isothermality (BIO-2, BIO-3) and many of the measured traits (e.g. flowering time, grain weight, flag leaf length) was observed on chromosome 4H (chr4H:619395701-621819033) where a flowering regulating gene *FT-like*, two *NAC* genes, the *COR27* gene, the stress induced *DIR* gene and more were identified (Fig. 2d). All these genes have a known role in tolerance to abiotic stress^38–41^ and are tightly linked within the same HB. Another example from chromosome 5H (chr5H:562546210-563321615), is associated with precipitation (BIO-12) and isothermality (BIO-3) and numerous traits (Fig. S7) including a flowering regulating gene (*CLAVATA*) was found next to *BTB/MATH*^42^, and gene which confers tolerance to abiotic stress^43^. All these genes are within a distance of circa 100Kbp. The high genomic proximity among these genes and many other stress-related genes (Table S3) can facilitate a synchronized and enhanced response to environmental stress including drought and heat.

### Adaptive risk assessment enables to prioritize areas for conservation and management

Climate change is expected to pose a significant challenge to populations failing to cope with the pace of environmental shifts. To estimate the potential maladaptive risk to the projected climate (2061-2080 years) among wild barley populations under mild (SSP1-2.6) and severe (SSP5-8.5) socioeconomic scenarios, a genetic offset analysis was conducted using the genomic composition of each population. For each scenario, three different climatic models^44^ (MPI-ESM1-2-HR, UKESM1-0-LL, and IPSL-CM6A-LR) were averaged and used to improve the robustness of the predictions. A spatial model was constructed within a gradient-forest framework using 14,564 SNPs from the identified regions harboring adaptive genes. The SNP dataset was used as the response matrix, and the 7 bioclimatic factors and 8 principal coordinates of neighbor matrices (PCNM) factors as predictors (“adaptive model”). Based on the models, the main bioclimatic factor risking populations persistence is annual precipitation (BIO-12) with an increased impact on the allelic composition at regions with rainfed with less than 250mm or above 750mm which marks the transition to the desert and high altitudes habitats, respectively (Fig. 3c). The next factor affecting the projected adaptive allelic composition is PCNM4 which corresponds to increased demographic isolation (isolation by distance) between populations. The third factor is isothermality (BIO-3) which reflects the day-to-night oscillations in temperatures that have a strong effect on development (growth degrees days), thus increased diurnal temperature variability is predicted to significantly constraint adaptation to future climate (Fig. 3a). To validate these results, a neutral model was constructed to reflect the proportion of variation explained merely by demographic processes and population structure. The model was calculated using a set of 100K SNPs randomly sampled across the genome. Overall, the neutral model explained less variation (mean *R^2^* = 0.27) than the adaptive model (mean *R^2^* = 0.69).

**Figure 3:**
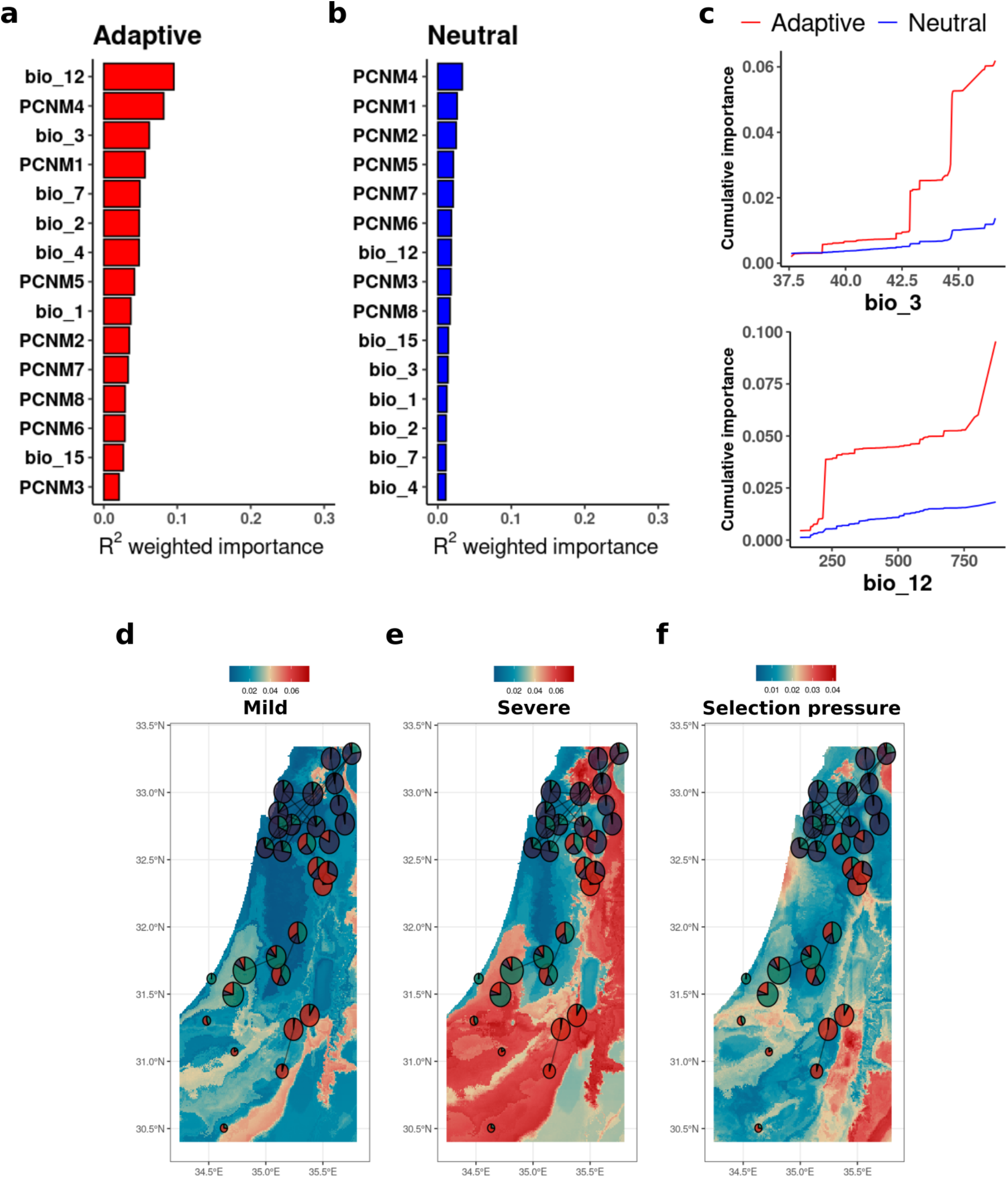
Predicting adaptive potential to climate change in wild barley populations. **a)** Relative contribution of the bioclimatic (BIOs) and spatial (PCNM) factors to the adaptive model. **b)** Relative contribution of the same factors to the neutral model. **c)** Cumulative importance of bioclimatic factors BIO-3 (top) and BIO-12 (bottom in the adaptive (red) and neutral (blue) models. **d)** Genetic offset analysis under mild scenario (SSP1-2.6)**. e)** Genetic offset analysis under severe scenario (SSP5-8.5). **f)** Residuals of the Procrustes rotation for the predicted allelic compositions by neutral and adaptive models. The color gradients from blue to red indicate regions under varying levels of selection pressure across wild barley populations. In all three panels (d-f), pie charts illustrate genetic clusters, with the size reflecting Tajima’s D score (smaller pie correspond to lower score), and lines between population pairs represent gene flow estimated via an isolation-by-distance approach.

To highlight habitats where wild barley populations are under selective pressure, a spatial comparison was conducted using Procrustes superimposition between the allelic composition predictions of adaptive and neutral models. Overall, extensive difference between models was observed for desert populations indicating a stronger selective pressure which is gradually declining towards the coastal and northern regions (Fig. 3f).

Next, the genetic offset was calculated as a Euclidean distance between present and projected future adaptive allelic compositions to assess the potential maladaptive risk. Geographically, higher maladaptive risk is expected mainly in isolated desert populations under mild scenario of climate change (SSP1-2.6). However, under the severe scenario (SSP5-8.5), risk is predicted to expand dramatically across the Levant, primarily in the desert (south) and along the Jordan valley (east). The observed difference between scenarios emphasizes the impact of policy and management on maladaptive risk. Significantly strong risk is also predicted in the southern coastal area and eastern Galilee in the north, where changes in temperature stability and precipitation may be too fast or excessive to enable local adaptation (Fig. 3d,e). At these regions, maintenance of ecological corridors, transplanting, and other management practices are specifically urged and expected to have the strongest benefit.

In several habitats, mainly along the northern coastal region of the Mediterranean Sea and at high elevations along the mountains ridge, a low maladaptive risk score was obtained (0.005-0.02) indicating an improvement of adaptation and suitability. These habitats are expected to obtain enough precipitation in order to provide refugia and enable the persistence of isolated populations (Fig. 3d,e). Interestingly, in some cases considerable risk was observed also in populations with high overall genomic diversity indicating that genetic variation alone is not a predictor for the adaptive potential (Fig. 3d,e). Finally, the results of the genetic offset model and predictions of suitability obtained by the SDM were compared using a Mantel test indicating a significant correspondence between ecological and evolutionary models (*r_M_* = 0.71; *p* = 0.001). Nevertheless, the genetic offset analysis was able to identify populations at high risk (e.g. southern coast and eastern Galilee) that were neglected by the species distribution model, recapitulating the importance of integrating genomic data into nature risk assessment models.

## Conclusions

Climate is changing at an accelerating pace and postulates a significant challenge for ecological systems and biodiversity. Our study provides important insights into the adaptive response of natural populations to the rapid climatic turnovers and the potential genetic mechanisms that regulate such response. First, our ecological-genetic sampling design enabled to overcome the masking effect imposed by the correlation between environmental and geographic distances that was a major constraint in previous studies^45–47^. Here, we were able to reduce the demographic effect and identify adaptive regions and candidate genes despite the high inbreeding and isolation by distance effects (Fig. 1c,d,h, 2c,d). The identified adaptive genetic variation was largely concentrated in haplotype blocks architecture, which enables to enhance the pleiotropic response to environmental change, supporting theoretical predictions^8–10^ and recent empirical studies^48^. Our results further established TEs as a responsive mechanism to enhance genetic variation specifically around adaptive haplotype blocks (Fig. 1g). TEs activation in response to environmental stress was shown to regulate recombination landscape and gene expression through methylation. Thus, the accumulation of TEs can mediate a rapid adaptive response to environmental changes.

The impact of projected climate change on wild barley populations in the southern Levant is mainly attributed to shifts in precipitation which was identified as the main limiting factor for the sustainability of populations (Fig. 3a). The models predict that the limited rainfall will funnel populations to creeks along the drainage basin or to higher elevations. This mode of dispersion will increase isolation among populations (Fig. 2a, 3d,e) enforcing them to sustain in climatic refugia, thus increasing the risk of maladaptation and potentially accelerating speciation and endemism^49^. Previous work has shown that migration in this region occurs mainly from north to south^33^. Assuming lower adaptive potential to drought and heat in the north, risk is expected to escalate with raising temperatures and reduction in precipitation. Moreover, the adaptive response of populations to projected climate is mainly reflected with a shift to shorter life cycle and early flowering, recapitulating the adoption of an escape strategy. This trend has been shown also in other species^50^ and is expected to broadly impact the entire ecological system including grazing fauna, grain-feeding insects and mammals, and their predators. Thus, future strategic planning of infrastructure development must consider the projected distribution of wild barley and likely other herbaceous species in the Levant to reduce risk of maladaptation and extinction of populations.

## Materials and methods

### Sampling design and plant material

The studied area lays within longitude 34-36 and latitude 29.3-33.5. To obtain a wide, yet balanced representation of the environmental and genetic variation of wild barley populations, sampling sites were selected based on climatic data obtained from the Israeli Meteorological Services (IMS), available eco-geographic raster layers, and available genetic information from the Barley1K collection^30^. The studied area was divided into six polygons. Three polygons were defined based on the assignment of populations from the Barley1K collection to ecotypes (>85% assignment), and three intermediate polygons between ecotypes regions were defined as ecotones. Within each polygon, five sampling sites were selected based on the environmental conditions, namely highest and lowest average annual temperature, lowest precipitation, highest precipitation variation between years, and one site with bright-rendzina soil type that can be found across all polygons (Fig. S1). The exact sampling locations were selected to maximize the representation of environmental variation and minimize anthropogenic disturbance. At each site, mature spikes were sampled from 10 plants that are separated by at least 10m from each other and geographical coordinates were recorded. Thus, 300 individual plants were sampled across 30 different sites. One seed from each plant was germinated and grown under controlled room conditions and was self-crossed to reduce potential heterozygosity. This procedure (single-seed-descent and selfing) was repeated three times to obtain a fixed accession from each plant.

### Climate data

In all models for present and future conditions we used the bioclimatic variables obtained from the WorldClim v2.1 database^33^. Non-relevant variables for this region were excluded and multicollinearity was controlled using a maximum correlation coefficient of 0.8 between bioclimatic variables. The remaining seven bioclimatic variables were BIO-1 (annual mean temperature), BIO-2 (mean diurnal range), BIO-3 (isothermality), BIO-4 (temperature seasonality), BIO-7 (temperature annual range), BIO-12 (annual precipitation), and BIO-15 (precipitation seasonality). Future climatic variables were obtained following a conservative approach by averaging three climate CMIP6 models^44^: MPI-ESM1-2-HR, UKESM1-0-LL, and IPSL-CM6A-LR, for 2061-2080 years under a mild (SSP1-2.6) and severe (SSP5-8.5) socioeconomics pathways. Data was downloaded from the WorldClim v2.1 database using the ‘geodata’ R package^51^ and processed with the ‘raster’ package^52^.

### DNA extraction, library preparation and sequencing

Genomic DNA was extracted from a leaf of a single young seedling using the CTAB buffer (100mM Tris HCl pH 8.0, 20mM EDTA pH 8.0, 1.4M NaCl, and 2% CTAB, 0.5% Sodium bisulfite), followed by Phenol/Chloroform purification protocol. Genomic DNA obtained from each sample was sheared to an average size of 350bp using Covaris E220. Sequencing libraries were prepared using the NEBNext® UltraTM II End Repair/dA-Tailing Module (NEB - E7546) followed by the NEBNext® UltraTM II Ligation Module (NEB - E7595). Samples were multiplexed with the Illumina-TruSeq DNA UD Indexes (96 indexes, 96 samples) (Illumina - #20022370), and post-ligation cleanup was done with X0.8 volume of homemade size-selection Magnetic-Beads (MB) washed twice with 80% ethanol. Libraries were amplified using KAPA HiFi HotStart ReadyMix (2X) (KAPABIOSYSTEMS - KK2600) and 10μM of Illumina compatible primers. Finally, each library was quantified using the QubitTM dsDNA HS Assay Kit (Invitrogen - Q32854) and qualified using high-sensitivity D1000 ScreenTape® (Agilent) to a mean size of 550bp. Pooling, titration, and sequencing on the NovaSeq 6000 system using the S4 flow cell with 2×150bp cycles were done at the Roy J. Carver Biotechnology Center, University of Illinois at Urbana-Champaign. Whole genome sequencing was targeted to at least 5X coverage for each accession and additional sequencing was performed for samples that did not generate adequate coverage.

### Variant calling

Raw reads were evaluated for quality and GC content using FastQC software v0.11.5^53^ and adapters, and low-quality data was trimmed using the fastp software^54^. Cleaned reads were aligned to the Morex reference genome v2^55^ using using BWA-MEM2 v2.2.1^56^ duplicated reads were marked using the gatk v4.1.9 and bam files were indexed with samtools v1.9^57^. Variant discovery analysis was conducted with HaplotypeCaller as implemented in gatk4^58^. In gatk, gvcf files were produced for each sample using the HaplotypeCaller tool and combined to a single data set with GenomicsDBImport and GenotypeGVCFs.

The obtained variants were filtered following the recommended hard filtering approach in gatk4 and based on the distribution of density plots generated for each statistic (Fig. S8). During filtering, only SNP data was kept and filtered for QualByDepth < 20, MappingQuality < 50, FisherStrand > 70, ReadDepth < 1000 and > 2000, QUAL < 140, StrandOddRatio > 2, and MappingQualityRankSum < −2.5. In addition, heterozygous sites were set as missing data, minimum minor allele frequency was set to 5%, maximum missing data was 10%, and the minimum number of reads to call an allele was 2.

All data processing and variant calling were conducted on the Amazon Web Services (AWS) system by splitting the pipeline into two phases. The first phase included the quality checks, trimming, alignment and bam files processing and generation of gvcf files; all these steps were performed for each sample independently, thus a total of 300 instances were deployed with 96 cores per instance. Once complete, all data was passed into an elastic file system. The second phase included the development of a genomic database and genotyping across all samples. To reduce the load, the gvcf files were divided into 1000 segments per chromosome, producing a total of 7,000 segments which were processed on 5,000 spot instances to reduce costs. Final vcf files were concatenated and filtered locally. For more details and code see https://github.com/hubner-lab.

### Phenotyping

Phenotyping was conducted in a common garden setup along three years in an insect-proof net-house located at the Hula valley, Israel (33°09’08.4”N 35°37’15.8”E). Before each growing season, seeds were stored in paper bags and oven-dried (30°C) for 14 days. After drying, seeds were dipped in 4% sodium hypochlorite solution for fungal prevention and were sown in mid-October in germination papers, watered, and put in dark-cold (4°C) conditions for 10 days for vernalization and dormancy break. After this treatment, germinating seedlings were transferred into 5-liter pots with commercial soil mix (“Ram 158”, Tuff Merom Golan Ltd, Merom Golan, Israel) and were grown under full irrigation for the rest of the growing season.

The common garden experiment was conducted in a randomized block design with 4 replicates for each genotype (Fig. S9). Phenotype data was collected during the growing seasons following the Zadoks scale definitions for development and phenology^58^. To facilitate this, data collection was performed daily using the FieldBook application v 5.0.7^60^. The dates at which each plant attained a developmental stage were recorded and subsequently converted into the number of days between each stage. To account for variations in developmental progress among genotypes throughout the growing season, time-based measurements were converted to Growing Degree Days (GDD) with a base temperature of 0°C. To achieve this, temperature data was sourced from the official meteorological service, specifically from the Kfar Blum station (no. 8473).

### Species distribution modeling

To quantify habitat suitability for wild barley across our study area, and to predict its spatial distribution, we fitted a species distribution model (SDM) using the random-forest algorithm^61^. We restricted our analyses to wild barley samples which occurred in sites with more than 100 mm of rainfall per year, resulting in 3281 presence points. Accordingly, we selected at random a sample of 3281 background points to reflect the distribution of environmental conditions in the entire region. We divided the data into training and testing sets based on an 80%-20% split, ensuring that each set had a nearly equal number of presence and background points. We extracted the values of seven environmental variables (distance to nearest river, mean diurnal temperature range, precipitation seasonality, precipitation of the wettest quarter, elevation above sea level, cation exchange capacity of the soil, and soil pH index measured in water solution) from their respective raster layers, in all presence and background points and training and testing sets. Using the values extracted from the training set, we fitted a random-forest model on the training data, with the dependent variable being the presence or absence of wild barley in the site (a binary variable). We then predicted the habitat suitability for wild barley across the entire study area by running the random-forest model on the full set of raster layers of environmental variables. Habitat suitability is a numeric value between 0 (low suitability) and 1 (high suitability). To evaluate model performance, we re-ran the model using the set-aside testing dataset and calculated its predictive accuracy according to the area under the curve (AUC) of the receiver operator curve (ROC).

To predict future distribution of wild barley, we re-ran the random-forest model developed using current environmental conditions, but this time using future conditions under two climatic scenarios (RCP45 and RCP85), for two years, 2050 and 2070. We generated predictions of habitat suitability for wild barley across the entire study area under each combination of year and scenario.

### Population structure Analyses

Population stratification analyses were conducted using called variants. To define population structure, variants were pruned to avoid the effect of linkage disequilibrium (LD) on population structure inference. The SNP dataset was LD-pruned with plink1.9 software^62^ with a minimum LD cut-off of r^2^ = 0.1, window size 100kb, and step size 20kb before conducting the PCA. Population structure was assessed using the sNMF function from LEA package^36^ with the number of ancestry populations (K) estimated based on principal component analysis (PCA) and cross-entropy test for K equals 3, 4, and 5 (Fig. S10).

The isolation by distance (IBD) between pairs of populations was determined using a Mantel test implemented in vegan R package^37^ between genetic distance and geographic distance. Genetic distance was calculated from the Q-matrix derived from sNMF analysis as Jaccard distance and geographic distance was calculated as Vincenty distance using the sampling sites coordinates. Isolation by environment (IBE) was determined using a Mantel test between the environmental distance calculated from the seven bioclimatic factors (BIO-1, BIO-2, BIO-3, BIO-4, BIO-7, BIO-12, BIO-15) as Euclidian distance and geographic distance.

Population genetics statistics including the average nucleotide diversity (π), and the Tajima’s D neutrality test were performed by chromosome using vcftools^63^ and averaged across the genome.

### Genome environment association analyses

Genome-environment association (GEA) analyses were performed using three different approaches: the redundancy analysis (RDA) implemented in the vegan package^37^, a mixed linear model implemented in the program EMMAX^35^, and a latent factor mixed model (LFMM) implemented in the LEA package^36^. Prior to the analysis, the missing variant calls were imputed based on the admixture coefficients (K=3) calculated with the ‘imputè function in the LEA package^36^. To improve the signal in genome-environment association (GEA) analysis, we calculated identity by state metric for each pair of individuals with plink software^60^ and filtered one sample from each pair which had identity by state higher than 0.99. After removing individuals, additional SNP filtering was applied based on minimum allele frequency 5% and maximum missing variants 10%. In EMMAX and RDA analyses, population structure was corrected using the first 3 principal components derived from a PCA as covariates. Additionaly, in EMMAX analysis a kinship matrix was used to correct relatedness as random effect. In LFMM, we used 15 latent factors to correct for population structure based on PCA and the cross-entropy criterion from sNMF analysis. To increase confidence in loci associated with environmental factors, the results of the different GEA approaches were combined (Fig. S4). Across all GEA analyses, significance was determined based on Bonferroni correction at 1.8×10^-9^.

### Genome-wide association study

Genome-wide association studies (GWAS) were performed for the imputed SNP dataset using the EMMAX software. To guarantee that imputation does not introduce bias, the analysis was conducted for few traits without imputation and the results were consisted. For correction of population structure the first 3 principal components were used and genetic relatedness was corrected based on a Balding-Nichols (BN) kinship matrix^35^.

Phenotype data was converted to BLUP scores using a mixed linear model implemented in the lme4 package^64^. Due to the stringent Bonferroni correction, a high false negative rate was observed, thus significant SNPs were determined at 1×10^-6^.

### Identifying haplotype blocks and candidate genes

Genome-wide haplotype blocks were identified using PLINK software v1.90^62^ after thinning the dataset and keeping a representative SNP per 1,000 bp leaving 3M variants. The data was further filtered for minor allele frequency < 0.2 to avoid the effect of rare alleles on the contingency of haplotypes. Linkage disequilibrium (LD) was calculated in a sliding window of 5Mbp and a minimum r² threshold of 0.2 was used to identify significant LD within windows. Haplotype blocks were identified using the --blocks command with a maximum block size set at 10Mbp. This approach enabled to focus on large, coherent haplotype blocks while minimizing the inclusion of unrelated genomic regions. Following this procedure, overlaps between significant SNPs in the GEA and GWAS analyses were used to identify adaptive haplotype blocks. Candidate genes were searched only in haplotype blocks that were supported by all GEA analyses and where associations with at least two phenotypic traits were identified (6 blocks).

### Quantifying TEs insertions

Transposable elements (TEs) insertions were quantified in 40 individuals from the most suitable and stressed environments based on the suitability scores derived from the SDM. To identify TEs insertions, we used the INSurVeyor software^65^ to inspect each accession alignment file (bam) separately. All identified insertions in each accession were aligned to the curated complete TREP database^66^ using blastn and excluding sequences with e-value higher than 1e^-5^. Number of TEs insertions were summarized for each accession across the entire genome, haplotype blocks, and in random regions of equal size sampled from the same chromosomes. To avoid bias, we excluded from the analyses the pericentromeric regions which were defined as 150Mbp around centromeres^55^.

### Genetic offset modeling

To model the genomic offset among populations we used a gradient forest analysis as implemented in the gradientForest package^67^. Two models were used to evaluate the vulnerability of populations to climate change. First, a ‘neutral’ model was trained using 100,000 randomly sampled SNPs across the entire genome. Next, an ‘adaptive’ model was trained using adaptive SNPs identified by intersecting significant SNPs from the GAWS analyses and GEA analyses. Thus, 14,564 SNPs were used for building ‘adaptive’ model. The maximum number of splits was calculated by the formula *log_2_(0.368*n / 2)*, where *n* is the number of sampling sites. The correlation threshold was set to 0.5 and the number of bootstrapped trees to 1000. In each model, allelic composition across SNPs was set as the response variable, the 7 major bioclimatic variables were included as predictors in addition to 8 principal coordinates neighbor matrices (PCNM) components calculated with vegan R package^37^ to correct the effect of spatial heterogeneity and structure. To model the effect of current and future environmental conditions the present and future bio-climatic variables were used, respectively.

To identify the extent of stress along space using genetic models, we performed a PCA transformation on the predictions from each model (’adaptive’ and ‘neutral’). Subsequently, we applied Procrustes rotation to align the datasets in a comparable orientation. The residuals were then extracted to estimate the differences between the ‘neutral’ and ‘adaptive’ models, thus high similarity indicates lower stress.

To estimate the adaptive risk in populations, we employed a genetic offset approach. Initially, projected future climate data were used to predict the genomic composition that would be adaptive under future climatic conditions. Genetic offset was then calculated as the Euclidean distance between the current adaptive genomic composition and the predicted future adaptive genomic composition.

Mantel test implemented in vegan R package was used to test for correlation between suitability scores obtained from the SDM and the genetic offset predictions for mild and severe socioeconomic scenarios using 999 permutations.

## Author contributions

SH conceived the study, DS and GA established the collection, TM, NA and DF conducted the sequencing, variant calling and phenotyping, EVP, MBK, ABM, performed the spatial analyses, EVP and SH performed the genomic analyses, EVP, ABM, and SH wrote the manuscript with input from all co-authors. All authors read and approved the manuscript.

## Data availability

Sequence data generated in this study are available on European Nucleotide Archive (ENA) project PRJEB79623, phenotype data are available through dryad XXX. Code and scripts are available through https://github.com/hubner-lab.

